# Should I shrink or should I flow? – body size adjustment to thermo-oxygenic niche

**DOI:** 10.1101/2020.01.14.905901

**Authors:** Aleksandra Walczyńska, Agnieszka Gudowska, Łukasz Sobczyk

## Abstract

Organisms adjust their size according to temperature and supposedly also respond to its negative covariate, oxygen. To what extent is size a response to temperature or oxygen? We analyzed the thermo-oxygenic niche for the community of 188 rotifer species. Evolution toward ranges of thermal tolerance occurred separately from evolution toward their optima. Body size was adjusted to both temperature and oxygen, but the cues for body size response differed; size was either driven by optimal temperatures or by the oxygen tolerance range. Animals are clearly separated into generalists or specialists, and their evolutionary body size adjustment is realized through differential responses to environmental factors. Oxygen is as important as temperature in the evolution of body size and ecological niche preference. An important conclusion from this study is that oxygen deprivation following global warming seems to be as problematic for the studied organisms as the temperature increase itself.

## Introduction

To understand the link between the performance of organisms and their environment is one of the *Grand Challenges* in organismal biology (Schwenk *et al*. 2009), especially in the context of abrupt climatic change (Allen *et al*. 2018). The general prediction is that body size decrease is a third universal response to global warming, especially in aquatic systems, following the geographic and phenological shifts in species distribution (Daufresne *et al*. 2009). Understanding the body size response to environmental factors is essential because this trait is unique (Kozłowski 2006). It can be perceived mutually as a morphological, physiological and life history trait, interconnecting the fields of ecology, evolution, and physiology. Despite the fundamental role of body size response, the key question of how animals become the size they are still awaits a satisfactory answer (Callier & Nijhout 2014). What kind of negative changes should we expect in the communities exposed to warming? Can we prevent at least some of these negative changes? Which actions should we undertake? What organisms do we save first? These urgent questions are not trivial for many reasons; however, the fundamental question of how organisms interact with their thermal environment remains unanswered. The crucial role of body size is that this trait is the major target of selective response on the organismal level (Kozłowski 2000), and such individual response affects the whole community through the plethora of possible ways including trophic interactions, dispersal abilities, habitat exploitation, nutrient cycling and others (Hildrew *et al*. 2007).

Ambient temperature is the most influential variable that shapes organismal strategies, from physical effects, through evolutionary influence, to ecological interactions (Schmidt-Nielsen 1990; Willmer *et al*. 2000; Begon *et al*. 2006). All life forms are equipped with mechanisms to detect and react to changing temperatures. Body size adjustment is an example of such a reaction. It is a phenomenon observed both genetically and phenotypically and is an assumed driver for interspecific Bergmann’s rule (Bergmann 1847) and the intraspecific temperature-size rule (Atkinson 1994). Body sizes changed with seasonal variations in temperature according to the rule “when it’s hot, shrink” in both small-scale (Kiełbasa *et al*. 2014) and large-scale studies (Horne *et al*. 2016; Horne *et al*. 2017). The empirical evidence for the direction of thermally induced body size changes on both the genetic and phenotypic levels in the same species is scarce, and it provides an ambiguous view of either positive covariance, as in *Brachionus plicatilis* rotifer (Walczyńska *et al*. 2017) or the negative one, as for the case of *Drosophila pseudoobscura* (Taylor *et al*. 2015).

Most other environmental variables are at least partly positively correlated with temperature. Oxygen is an exception because it is negatively correlated with temperature (Wetzel 2001 for aquatic systems). Oxygen stress often occurs at higher temperatures, because the energetic demands of an organism grow faster than the oxygen supply (Verberk *et al*. 2011). This phenomenon is especially important for organisms that inhabit aquatic environments (Forster *et al*. 2012; Horne *et al*. 2015; Horne *et al*.2017). It is caused by the fact that breathing under water is much more challenging than in air because of the much slower oxygen diffusion in the former system (Verberk *et al*. 2011). Oxygen stress at higher temperatures has been suggested to cause the cell size to decrease to enhance the efficiency of oxygen transport to the mitochondria (Woods 1999), and as a result, the whole body shrinks (Atkinson *et al*. 2006). Animals anticipate oxygen deficiency when experiencing a temperature increase (Walczyńska *et al*. 2015). Thus, they face the ecophysiological dilemma of either accepting this challenge and adjusting their cell (and body) size to overcome the reduced aerobic metabolism efficiency or behaviorally adapting to more favorable conditions. Those able to decrease their size will respond until the conditions are too stressful, while those lacking the ability to respond are immediately at risk when escape is not possible. To understand whether and how organisms respond to change, we first need to identify the exact environmental response cues and initiation mechanisms. Including oxygen concentrations as a covarying parameter in the studies on size-to-temperature response facilitates the interpretation of the results (Kiełbasa *et al*. 2014; Walczyńska & Sobczyk 2017). A main question has emerged: what exactly drives body size responses?

Here, we used an exceptional field model to study evolutionary outputs that could not be experimentally tested in the laboratory. We scrutinized data published 30 years ago in the new context of thermo-oxygenic niche construction within an aquatic community. The original data were provided by Bērzinš and Pejler, who reported the results of planktic, periphytic and benthic rotifer sampling from approximately 600 different water bodies (lakes, ponds, rivers and mires) in Sweden for approximately 40 years. The authors related the occurrence of each species to temperature (Bērzinš & Pejler 1989b) and oxygen concentration (Bērzinš & Pejler 1989a), among other factors. The data, supplemented with information on species-specific body sizes, gave us a unique opportunity to study the interspecific size response to subtle aspects of the environment at the macroevolutionary scale, in the context of ecological niche.

The evolution of the thermal niche is still an ecological riddle; natural selection shapes the thermal optimum, breadth of tolerance range and performance limit, but the correlations between these traits remain unknown, preventing an understanding of the ecological diversity of life (Mongold *et al*. 2008). An extension of this issue is whether species with a specific thermal range specialize at the same level along other niche axes (Sheth & Angert 2014). To answer these questions, we analyzed how similar the preferences for different thermal and oxygenic conditions were within a community of rotifers. We also refer to an important demarcation line in the strategy of dealing with the environment, through a distinction between generalists, organisms that display the relatively high and flat performance across environments, and specialists, those that perform better in one type of environment than in all the others (Levins 1968). We examined how species-specific standard body size was affected by the joint thermal and oxygenic conditions of living. To find the subtle environmental cues for the possible size differences we preceded with the multivariate analyses to reveal how the species preferences described by the temperature and oxygen optima, ranges and tolerance limits were interrelated at the community level. We predicted the species body size to be affected by both the temperature and oxygen.

We provide the first evidence of sharing the two-dimensional ecological niche, and of interspecific evolutionary body size response to this niche, for the large community of aquatic organisms.

## Material and methods

### Preparation of the dataset

Each rotifer species was characterized by eight parameters describing its environmental preferences, namely, minimum/maximum/optimum/range of temperature/oxygen concentration in the living habitat. We obtained data on these species-specific parameters from two publications: Bērzinš and Pejler (1989a)(minimum, maximum, optimum and range of tolerance to oxygen concentration) and Bērzinš and Pejler (1989b) (the same parameters describing tolerance to temperature). We interpreted the values from the figures using millimeter paper, with an accuracy of 0.5 mg/L for O_2_ and 1 °C for temperature. In both cases, the tolerance range was calculated as *max - min*, while the optimal value was assumed to be the value with the maximal abundance, as presented in the respective original figures. To compare the variability within the environmental variables examined, we estimated the coefficient of quartile variation (CQV), calculating (*Q_3_-Q_1_*)/(*Q_3_*+*Q_1_*), where *Q_1_* is a first quartile and *Q_3_* is a third quartile, for each variable separately.

The authors of the original articles did not provide data on the sizes of the species, but we were interested in the relative differences in rotifer species-specific body size rather than their local adaptations. Thus, we analyzed the association between the species standard body length (μm) of rotifers, collected from the available databases, including Bielańska-Grajner *et al*. (2013), Bielańska-Grajner *et al*. (2015), Kreutz and Foissner (2006), Segers (1995), Segers and Shiel (2005) and the database from the website of the National Institute for Environmental Studies (http://www.nies.go.jp/), by searching for the species Latin name in a website browser. We standardized the length of the body by using the values provided for fixed, nonstretched individuals and excluding the length of toes and other appendages. In the cases when a size range was provided, we calculated the mean value. When more than one dataset was provided, especially for cited websites, we calculated the mean for all the subsources. The dataset for the 188 rotifer species, with the species-specific body lengths and sources of information, are provided in the supplementary materials (Table S1).

### PCA analysis

To determine how the parameters describing the thermal and oxygenic preferences grouped at the interspecific level, we conducted the principal component analysis (PCA), with rows representing the species and columns represented by eight environmental parameters. We log-transformed and standardized the data to provide the correlation matrix and we conducted the analysis in CANOCO 5.0 (Ter Braak & Šmilauer 2012).

### Phylogenetic analysis

We performed all phylogenetic analyses in the R computational environment (v3.4.0) (R Core Team 2017). We obtained the phylogeny of 188 rotifer species from the open tree of life (Hinchliff *et al*. 2015) and ‘rotl’ package v3.0.3 (Michonneau *et al*. 2016). Because the branch lengths were not available, we automatically estimated them using a method proposed by Grafen (1989) and the ‘ape’ package v4.1 (Paradis *et al*. 2004). All the path lengths from the root to the tips were equal. The tree contained polytomies, where more than two branches descended from a single node. Simulation studies have generally found that independent contrasts and phylogenetically generalized least squares (PGLS) are fairly robust to errors in both phylogenetic topology and branch length (Symonds & Blomberg 2014). We tested the relationship between species body size (length, after natural logarithm transformation) and species-specific environmental characteristics in a phylogenetic comparative model. We applied PGLS using the gls() function in ‘nlme’ in the caper package (https://CRAN.R-project.org/package=caper) considering body size as a response variable and four environmental parameters (ranges and optima) as explanatory variables. In the model, we assumed Brownian motion (BM) evolution, which is the most commonly assumed model of phenotypic evolution by comparative phylogenetic methods (Revell 2010; Lajeunesse & Fox 2015). BM evolution treated random genetic drift as a primary process resulting in the loss of similarity from ancestral characteristics (Martins & Garland 1991). Traits, e.g., body size and morphology, exhibit strong phylogenetic signals and most likely evolve by gradual changes over time according to the BM model of evolution (Symonds & Blomberg 2014). We assumed the correlation structure based on Pagel’s λ (Pagel 1999) fixed at 1.

## Results

### Thermo-oxygenic niche of rotifer assemblage

PCA analysis showed three informative principal components (PCs) for data interpretation (eigenvalue > 1), explaining 37.47 % (PC1), 29.17 % (PC2) and 15.08 % (PC3) of the variance (Fig. 1A, B). The evolutionary adaptation may be studied as a process or a product (Mongold *et al*. 2008). Treating the community of rotifers as a gene pool, we actually observed a product of evolution (Kimura 1974) at the assemblage level; the evolution of generalists was affected by both temperature and oxygen, as explained by PC1 (horizontally oriented arrows for temperature and oxygen tolerance ranges in Fig. 1A and their high loadings in Fig. 1B), and was clearly separated from the evolution of specialists, as explained by PC2 (vertically oriented arrows for temperature and oxygen optima in Fig. 1A and their high loadings in Fig. 1B). The generally opposite position of the parameters representing temperature *vs*. oxygen in PC2 (Fig. 1A, C) reflects the importance of their natural negative correlation in evolutionary processes. PC3 shows the ecological force of minor importance that caused a non-uniformity in the pattern of breath of thermal tolerance and hypoxia tolerance: according to PC3 the relationship between Trange and O2min is positive, as compared to their negative link according to PC1 in Fig. 1B). According to the CQV analysis, the lowest variability was observed for T_max_, oxygen optimum (O_2opt_) and oxygen maximum (O_2max_), followed by optimum temperature (T_opt_), temperature range (T_range_), and oxygen tolerance range (O_2range_) (Fig. 3C).

**Fig. 1.**
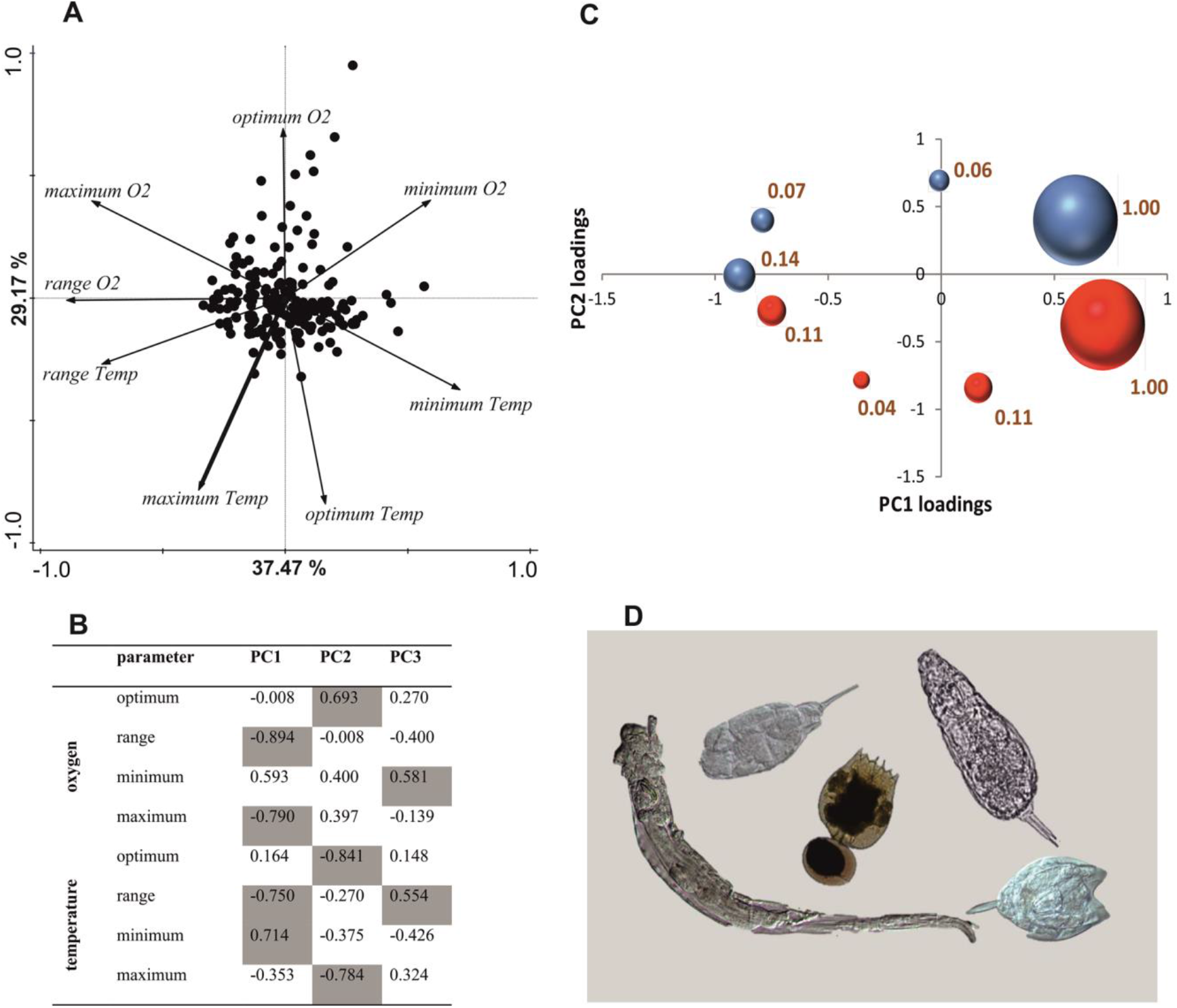
The evolution of the thermo-oxygenic niche in 188 rotifer species. The results of the PCA analysis: **A** – factor plane (PC1-PC2) projection (each point represents a species), **B** – parameter loadings for PC1-PC3 in a multivariate space, **C** – coefficient of quartile variation (CQV) of each parameter represented by bubble size on the PC1/PC2 plane. The parameters are in the same order as in **(A)**; temperature parameters are marked in red and oxygen parameters are in blue. The CQV value of each parameter is provided next to the respective bubble. Exemplary rotifer species are presented in **D**.

### Body size of species assembling the niche

We tested the body size relationship to the parameters representing the PC1 (ranges) and PC2 (optima). Phylogenetically corrected regression analyses based on data obtained from the open tree of life (Hinchliff *et al*. 2015) for 188 species (Fig. 2) revealed that rotifer body size increased with increasing tolerance to oxygen range (O_2range_, p < 0.01; adjusted R^2^ = 0.054), decreased with increasing optimal temperature (T_opt_, p < 0.01; adjusted R^2^ =0.048), and had no relationship with the remaining two parameters (p = 0.20 and adjusted R^2^ = 0.003 for T_range_ and p = 0.57 and adjusted R^2^ = - 0.004 for O_2opt_; Fig. 3). Body size evolved in response to both variables, temperature and oxygen, but the evolutionary cues for response were different; species were smaller when specializing to a high optimal temperature or to a narrow oxygen tolerance range (Fig. 3).

**Fig. 2.**
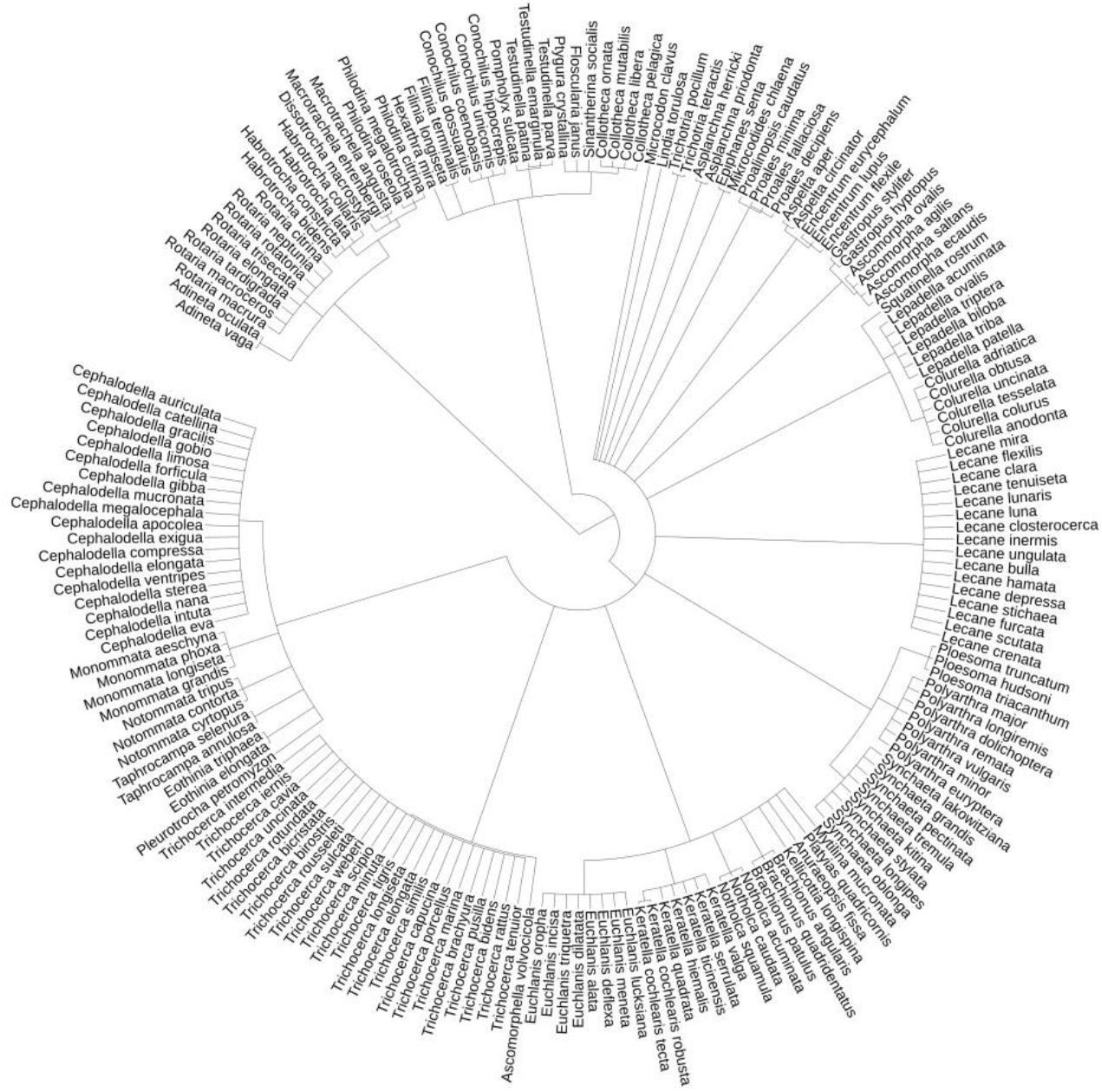
Phylogenetic tree for all rotifer species representing a dataset for those species with available phylogenetic data (N = 188). Data were collected from the tree of life web project. The branch lengths were arbitrarily estimated using the method proposed by Grafen (1989) and the ‘ape’ package v4.1 (Paradis *et al*. 2004).

## Discussion

We found that within the rotifer community that involved 188 species of different ecology (planktic, periphytic and benthic), representing various aquatic habitats (i) species-specific thermal tolerance range and oxygen tolerance range evolved in the same direction, (ii) optima for temperature and oxygen evolved in opposite directions. These results raise an intriguing question regarding the possible different physiological mechanisms behind the selective forces of adaptation to tolerance ranges *vs*. specialization to specific optima. They also mean that, in general, a species characterized by the wide range of thermal tolerance should be expected to have the wide range of tolerance to oxygen availability as well.

The analysis of body size relationship with eight environmental variables which describe the thermo-oxygenic niche showed that, at the interspecific level, body size decreased in response to both temperature and oxygen, but in different ways; the target of size response was optimum in the case of temperature and the tolerance range in the case of oxygen.

The interspecific variability in each of the eight parameters shows that oxygen changes will affect the organisms similarly to temperature changes as a consequence of the current climate changes. Araújo *et al*. (2013) estimated the variance in cold tolerance (CT_min_) *vs*. heat tolerance (CT_max_) in different groups of ectotherms. They found that the cold tolerance variance was almost twice as high as the heat tolerance variance. Their interpretation of the results was that ectotherms are more vulnerable to an increase in maximum temperatures than in minimum temperatures; tolerance to cold is labile and subject to natural selection, whereas tolerance to heat is physiologically conserved. In this respect, our results are in agreement; the tolerance to cold was more variable than the tolerance to heat (7.82 *vs*. 5.98 of variance for T_min_ and T_max_, respectively). A similar comparison between oxygen and temperature requires the use of a scale-independent parameter, such as CQV. Provided that the reasoning of Araújo et al. is correct, the rotifer community is most vulnerable to changes in these variables, while the least conserved are Tmin and low oxygen (hypoxia!) tolerance (O_2min_). The result obtained for species-specific oxygen optimum and maximum seems to be especially important considering the global ocean is undoubtedly warming (Cheng *et al*. 2019), and its oxygen concentration is decreasing through different synergistic mechanisms (Breitburg *et al*. 2018). Our CQV analysis showed that rotifer community appeared sensitive not only to potential changes in upper thermal limits (T_max_) but similarly much to oxygen deprivation below the optimal concentrations at which the species perform best (O_2opt_; Fig. 1C). This result acts as a specific warning: aquatic ectotherms are potentially very vulnerable to climate warming because the successive absolute reduction in oxygen concentration in water may be too challenging for them to quickly adapt.

The exceptional feature of the data published thirty years earlier is that by tracking the members of the community in their natural habitat, we can actually observe the ecological, realized niche, which is a component of the fundamental niche remaining for usage in consequence of interactions with other organisms (Hutchinson 1957). Hence, it seems justified to claim that the here-studied rotifer community construct their ecological niche by detecting and responding to subtle environmental cues. Within their thermo-oxygenic niche, the parameters limiting the interspecific body size adjustment are maximal temperature, optimal oxygen and maximal oxygen concentration.

### Generalist-specialist continuum implications

The PCA showed that evolution toward ranges of temperature and oxygen tolerance occurred separately from evolution toward specialization for high performance at pick values of both parameters. This clear trade-off between evolution toward generalist and specialist strategies agree with current theoretical predictions (Levins 1968; Huey & Hertz 1984). As a consequence of distinguished patterns of adaptation to ranges of tolerance or specialization for certain optimal values, the simple observation of decreasing body size with increasing temperature within the thermo-oxygenic niche may be an outcome of two different physiological mechanisms leading to the same result: a large size at low temperatures and high relative oxygen availability and a small size at high temperatures and low relative oxygen availability (Fig. 4). This retorts to the matter whether decreasing in heat is indeed equivalent to increasing in cold (Walczynska *et al*. 2018). The former is driven by oxygen-limited aerobic metabolism (Woods 1999; Verberk *et al*. 2011) and may evolve along with a narrow tolerance to oxygen conditions (this study), while the latter may be driven by the limitation of proper genome maintenance in cold conditions (Xia 1995; Woods *et al*. 2003; Hessen *et al*. 2013) and/or the energetic costs of cell membrane maintenance (Szarski 1983) and may reflect specialization toward a low optimal temperature. The existence of a common explanation for body size-to-temperature observations was questionable (Angilletta & Dunham 2003; Angilletta Jr 2009). The dual causative mechanism we report here constitutes such a possible common, though complex, clarification.

### Global change implications

It is imperative to elucidate whether the cues species experience are clear enough to respond. The warming effect is not symmetrical, as the minimum temperature is rising faster than its maximum (Easterling *et al*. 1997), and species with preferences for low optimal temperatures or high optimal oxygen concentrations are challenged differently than species with preferences in the upper thermal range. High optimal oxygen concentration is distinctive in this regard because species are apparently specialized for specific O2opt by some mechanisms alternative to the body size adjustment at the interspecific level (Fig. 3); neither they would be able to escape when facing a large-scale process of oxygen deprivation, such as global warming. With regard to the questions posed at the beginning of this text, our result on the high conservativity regarding preferred oxygen levels has important implications for conservation strategies. We should immediately start by focusing on saving the ecosystems with the highest risk of a sharp decline in oxygen availability.

**Fig. 3.**
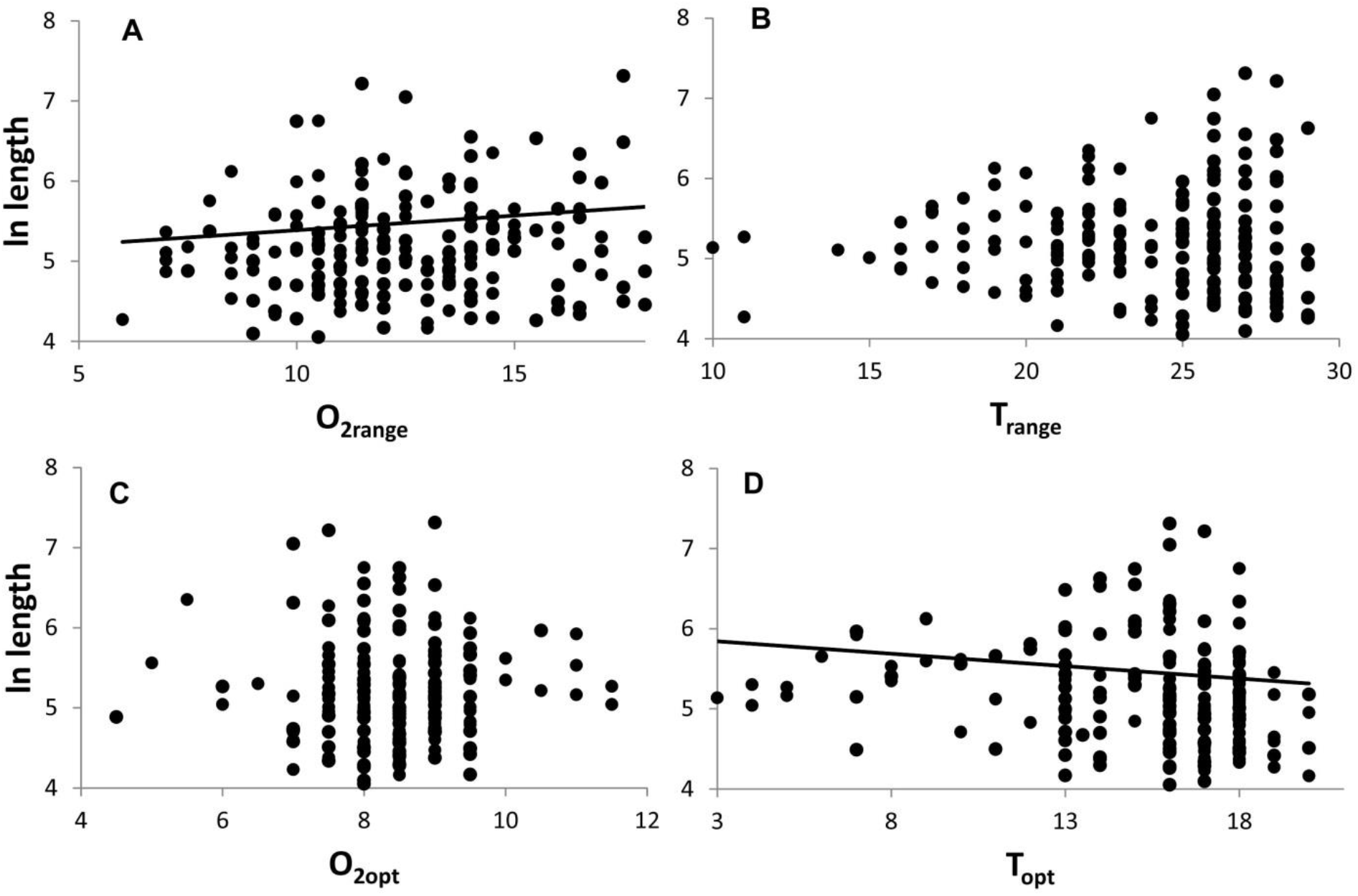
Body size evolution toward a thermo-oxygenic niche in 188 rotifer species. The simple regression estimates of the relationship of body length with four environmental parameters that drove the PCA: ranges (A, B) and optima (C, D) of oxygen concentration and temperature. Each point represents an original value for a given species, while the estimation was phylogenetically corrected. Significant relationships are shown with their linear estimations.

**Fig. 4.**
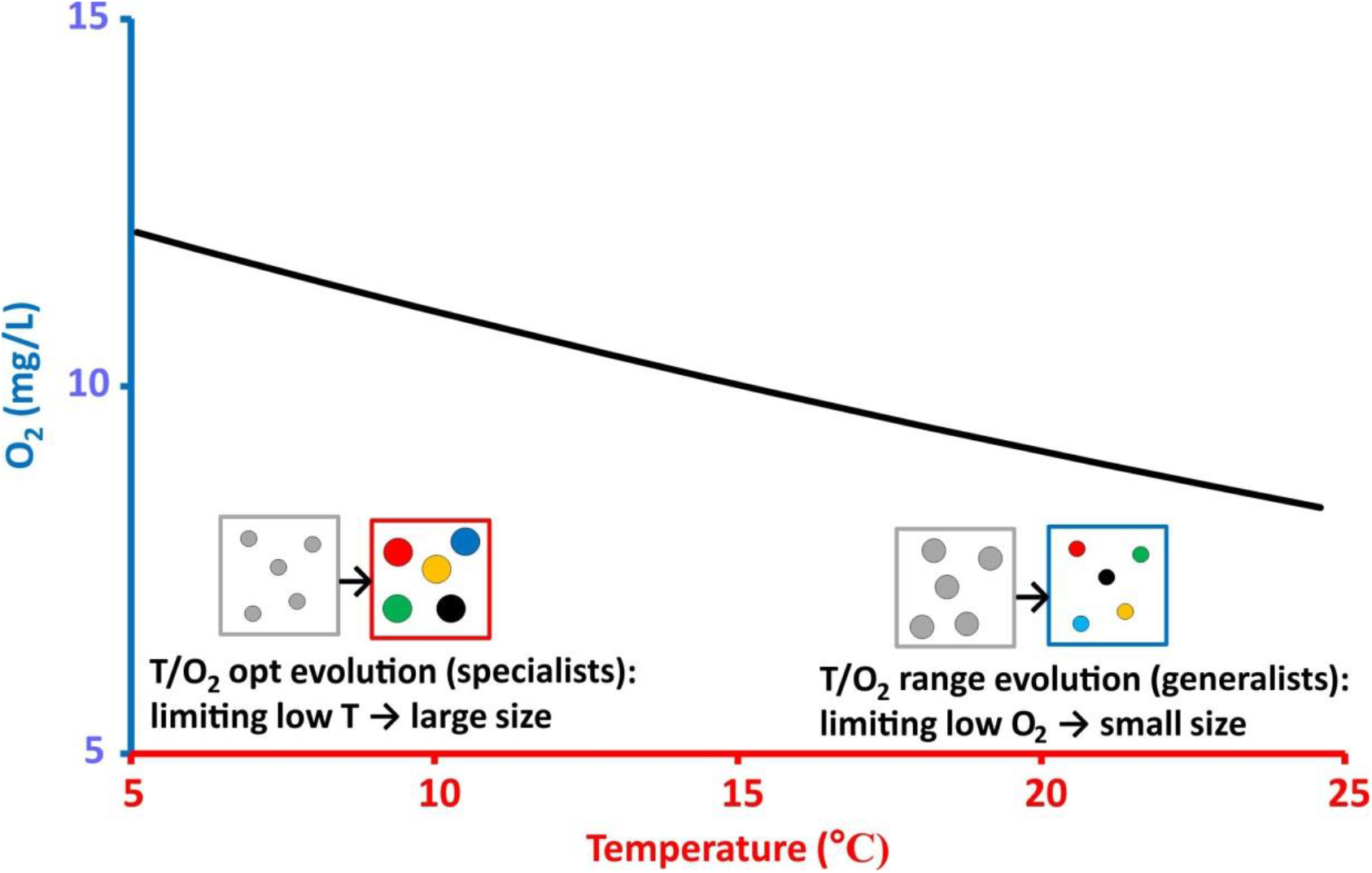
The thermo-oxygenic niche and its relationship to body size evolution in the assemblage of 188 rotifer species. The communities consist of large species at low temperature/high O_2_ conditions and small species at high temperature/low O_2_ conditions. This common observation may result from two compatible processes: low temperature may limit the small size, while low O_2_ availability may constrain the large size. We associated this pattern with the evolution toward the generalist or specialist strategy, which was clearly divided according to our PCA. The pattern of O_2_ decrease with increasing temperature (in water) is based on data from Wetzel (2001).

To conclude, in this study we show that the community of aquatic animals displays clear preferences within a thermo-oxygenic niche, which reflects in the species-specific body size response to subtle cues of both environmental variables studied. This is the first evidence of such a clear pattern of within-community response to two-dimensional ecological niche, that may act as a base for any ecological large-scale studies and models, especially regarding the consequences of global warming. Our strong message is that oxygen should be taken into account as a variable similarly important to temperature, if we aim to understand, or to counteract, the effects of climatic changes on communities.

## Acknowledgments

The authors are thankful to Ulf Bauchinger, Terézia Horváthová and Manuel Serra for the very helpful comments on the previous versions of this manuscript. This study would not be possible without the substantial work done long ago by Bruno Bērzinš and Birger Pejler.

