## Supplementary Table S1 for "Should I shrink or should I flow? – body size adjustment to thermo-oxygenic niche"

**Table S1.** The list of all the rotifer species analyzed, their species-specific body mass and the bibliographic source of body size information.

| No. | species | mean body length (um) | bibliographic source |
|---|---|---|---|
| 1 | *Adineta oculata* | 444 | Segers & Shiel 2005 |
| 2 | *Adineta vaga* | 455 | Bielańska-Grajner et al. 2013 |
| 3 | *Anuraeopsis fissa* | 99 | http://www.nies.go.jp; Ejsmont-Karabin et al. 2004 |
| 4 | *Ascomorpha agilis* | 145 | Ejsmont-Karabin et al. 2004 |
| 5 | *Ascomorpha ecaudis* | 165 | Ejsmont-Karabin et al. 2004 |
| 6 | *Ascomorpha ovalis* | 150 | Kreutz & Foissner 2006; Ejsmont-Karabin et al. 2004 |
| 7 | *Ascomorpha saltans* | 122 | Kreutz & Foissner 2006; Ejsmont-Karabin et al. 2004 |
| 8 | *Ascomorphella volvocicola* | 130 | Ejsmont-Karabin et al. 2004 |
| 9 | *Aspelta aper* | 263 | http://www.nies.go.jp ; Ejsmont-Karabin et al. 2004 |
| 10 | *Aspelta circinator* | 230 | Ejsmont-Karabin et al. 2004 |
| 11 | *Asplanchna herricki* | 854 | Ejsmont-Karabin et al. 2004 |
| 12 | *Asplanchna priodonta* | 756 | http://www.nies.go.jp; Ejsmont-Karabin et al. 2004 |
| 13 | *Brachionus angularis* | 121 | http://www.nies.go.jp; Ejsmont-Karabin et al. 2004 |
| 14 | *Brachionus patulus* | 110 | http://www.nies.go.jp |
| 15 | *Brachionus quadridentatus* | 234 | http://www.nies.go.jp; Ejsmont-Karabin et al. 2004 |
| 16 | *Cephalodella catellina* | 115 | http://www.nies.go.jp; Ejsmont-Karabin et al. 2004 |
| 17 | *Cephalodella apocolea* | 143 | http://www.nies.go.jp; Ejsmont-Karabin et al. 2004 |
| 18 | *Cephalodella auriculata* | 131 | http://www.nies.go.jp; Kreutz & Foissner 2006; Ejsmont-Karabin et al. 2004 |
| 19 | *Cephalodella compressa* | 142 | http://www.nies.go.jp |
| 20 | *Cephalodella elongata* | 110 | http://www.nies.go.jp |
| 21 | *Cephalodella eva* | 122 | http://www.nies.go.jp; Ejsmont-Karabin et al. 2004 |
| 22 | *Cephalodella exigua* | 89 | http://www.nies.go.jp; Ejsmont-Karabin et al. 2004 |
| 23 | *Cephalodella forficula* | 198 | http://www.nies.go.jp; Ejsmont-Karabin et al. 2004 |
| 24 | *Cephalodella gibba* | 217 | http://www.nies.go.jp; Kreutz&Foissner |
| 25 | *Cephalodella gracilis* | 107 | http://www.nies.go.jp; Ejsmont-Karabin et al. 2004 |
| 26 | *Cephalodella gobio* | 111 | http://www.nies.go.jp; Ejsmont-Karabin et al. 2004 |
| 27 | *Cephalodella intuta* | 86 | http://www.nies.go.jp |
| 28 | *Cephalodella limosa* | 111 | Ejsmont-Karabin et al. 2004 |
| 29 | *Cephalodella megalocephala* | 173 | http://www.nies.go.jp; Ejsmont-Karabin et al. 2004 |
| 30 | *Cephalodella mucronata* | 150 | http://www.nies.go.jp |
| 31 | *Cephalodella nana* | 80 | http://www.nies.go.jp; Ejsmont-Karabin et al. 2004 |
| 32 | *Cephalodella sterea* | 148 | http://www.nies.go.jp; Ejsmont-Karabin et al. 2004 |
| 33 | *Cephalodella ventripes* | 111 | http://www.nies.go.jp; Ejsmont-Karabin et al. 2004 |
| 34 | *Collotheca libera* | 97 | Ejsmont-Karabin et al. 2004 |
| 35 | *Collotheca mutabilis* | 182 | http://www.nies.go.jp |
| 36 | *Collotheca ornata* | 139 | http://www.nies.go.jp; Kreutz & Foissner 2006 |
| 37 | *Collotheca pelagica* | 159 | http://www.nies.go.jp; Ejsmont-Karabin et al. 2004 |
| 38 | *Colurella adriatica* | 90 | http://www.nies.go.jp; Ejsmont-Karabin et al. 2004 |
| 39 | *Colurella anodonta* | 60 | http://www.nies.go.jp |
| 40 | *Colurella colurus* | 83 | http://www.nies.go.jp; Ejsmont-Karabin et al. 2004 |
| 41 | *Colurella obtusa* | 73 | Kreutz & Foissner 2006; Ejsmont-Karabin et al. 2004 |
| 42 | *Colurella tesselata* | 58 | Ejsmont-Karabin et al. 2004 |
| 43 | *Colurella uncinata* | 81 | http://www.nies.go.jp; Ejsmont-Karabin et al. 2004 |
| 44 | *Conochilus dossuarius* | 310 | http://www.nies.go.jp; Ejsmont-Karabin et al. 2004 |

| | | | |
|---|---|---|---|
| 45 | *Conochilus coenobasis* | 275 | http://www.nies.go.jp |
| 46 | *Conochilus hippocrepis* | 565 | http://www.nies.go.jp |
| 47 | *Conochilus unicornis* | 285 | http://www.nies.go.jp |
| 48 | *Dissotrocha macrostyla* | 442 | Kreutz & Foissner 2006; Bielańska-Grajner et al. 2013 |
| 49 | *Encentrum eurycephalum* | 390 | Ejsmont-Karabin et al. 2004 |
| 50 | *Encentrum flexile* | 194 | http://www.nies.go.jp |
| 51 | *Encentrum lupus* | 175 | Ejsmont-Karabin et al. 2004 |
| 52 | *Eothinia elongata* | 388 | Ejsmont-Karabin et al. 2004 |
| 53 | *Eothinia triphaea* | 182 | http://www.nies.go.jp |
| 54 | *Epiphanes senta* | 458 | http://www.nies.go.jp; Ejsmont-Karabin et al. 2004 |
| 55 | *Euchlanis alata* | 262 | Ejsmont-Karabin et al. 2004 |
| 56 | *Euchlanis dilatata* | 228 | Ejsmont-Karabin et al. 2004 |
| 57 | *Euchlanis deflexa* | 290 | http://www.nies.go.jp |
| 58 | *Euchlanis lucksiana* | 186 | http://www.nies.go.jp |
| 59 | *Euchlanis incisa* | 224 | Ejsmont-Karabin et al. 2004 |
| 60 | *Euchlanis meneta* | 137 | Ejsmont-Karabin et al. 2004 |
| 61 | *Euchlanis oropha* | 169 | Ejsmont-Karabin et al. 2004 |
| 62 | *Euchlanis triquetra* | 431 | Ejsmont-Karabin et al. 2004 |
| 63 | *Filinia longiseta* | 168 | Ejsmont-Karabin et al. 2004 |
| 64 | *Filinia terminalis* | 168 | Ejsmont-Karabin et al. 2004 |
| 65 | *Floscularia janus* | 215 | Ejsmont-Karabin et al. 2004 |
| 66 | *Gastropus hyptopus* | 229 | http://www.nies.go.jp; Ejsmont-Karabin et al. 2004 |
| 67 | *Gastropus stylifer* | 172 | Ejsmont-Karabin et al. 2004 |
| 68 | *Habrotrocha bidens* | 411 | http://www.nies.go.jp |
| 69 | *Habrotrocha collaris* | 292 | http://www.nies.go.jp |
| 70 | *Habrotrocha constricta* | 287 | http://www.nies.go.jp |
| 71 | *Habrotrocha lata* | 196 | http://www.nies.go.jp |
| 72 | *Hexarthra mira* | 286 | http://www.nies.go.jp; Ejsmont-Karabin et al. 2004 |
| 73 | *Kellicottia longispina* | 687 | http://www.nies.go.jp; Ejsmont-Karabin et al. 2004 |
| 74 | *Keratella cochlearis robusta* | 210 | http://www.nies.go.jp |
| 75 | *Keratella cochlearis tecta* | 100 | http://www.nies.go.jp |
| 76 | *Keratella hiemalis* | 200 | Ejsmont-Karabin et al. 2004 |
| 77 | *Keratella quadrata* | 229 | Ejsmont-Karabin et al. 2004 |
| 78 | *Keratella serrulata* | 181 | Ejsmont-Karabin et al. 2004 |
| 79 | *Keratella ticinensis* | 132 | Ejsmont-Karabin et al. 2004 |
| 80 | *Keratella valga* | 194 | Ejsmont-Karabin et al. 2004 |
| 81 | *Lecane depressa* | 121 | http://www.nies.go.jp; Ejsmont-Karabin et al. 2004 |
| 82 | *Lecane bulla* | 142 | Kreutz & Foissner 2006; Ejsmont-Karabin et al. 2004 |
| 83 | *Lecane clara* | 99 | Segers 1995; Ejsmont-Karabin et al. 2004 |
| 84 | *Lecane closterocerca* | 71 | Ejsmont-Karabin et al. 2004 |
| 85 | *Lecane flexilis* | 73 | Ejsmont-Karabin et al. 2004 |
| 86 | *Lecane furcata* | 64 | http://www.nies.go.jp |
| 87 | *Lecane hamata* | 80 | http://www.nies.go.jp; Ejsmont-Karabin et al. 2004 |
| 88 | *Lecane inermis* | 91 | Kreutz & Foissner 2006; Segers 1995; Ejsmont-Karabin et al. 2004 |
| 89 | *Lecane luna* | 131 | http://www.nies.go.jp; Ejsmont-Karabin et al. 2004 |
| 90 | *Lecane lunaris* | 110 | http://www.nies.go.jp; Ejsmont-Karabin et al. 2004 |
| 91 | *Lecane crenata* | 110 | http://www.nies.go.jp |
| 92 | *Lecane mira* | 143 | http://www.nies.go.jp; Ejsmont-Karabin et al. 2004 |

| | | | |
|---|---|---|---|
| 93 | *Lecane scutata* | 72 | Ejsmont-Karabin et al. 2004 |
| 94 | *Lecane stichaea* | 90 | Ejsmont-Karabin et al. 2004 |
| 95 | *Lecane tenuiseta* | 69 | Ejsmont-Karabin et al. 2004 |
| 96 | *Lecane ungulata* | 216 | http://www.nies.go.jp; Ejsmont-Karabin et al. 2004 |
| 97 | *Lepadella acuminata* | 551 | Ejsmont-Karabin et al. 2004 |
| 98 | *Lepadella biloba* | 93 | http://www.nies.go.jp |
| 99 | *Lepadella ovalis* | 135 | Ejsmont-Karabin et al. 2004 |
| 100 | *Lepadella patella* | 86 | http://www.nies.go.jp; Ejsmont-Karabin et al. 2004 |
| 101 | *Lepadella triptera* | 72 | http://www.nies.go.jp; Ejsmont-Karabin et al. 2004 |
| 102 | *Lepadella triba* | 77 | http://www.nies.go.jp; Ejsmont-Karabin et al. 2004 |
| 103 | *Lindia torulosa* | 315 | Ejsmont-Karabin et al. 2004 |
| 104 | *Macrotrachela angusta* | 275 | http://www.nies.go.jp |
| 105 | *Macrotrachela ehrenbergi* | 312 | http://www.nies.go.jp |
| 106 | *Mikrocodides chlaena* | 169 | http://www.nies.go.jp |
| 107 | *Microcodon clavus* | 192 | Ejsmont-Karabin et al. 2004 |
| 108 | *Monommata aeschyna* | 150 | http://www.nies.go.jp |
| 109 | *Monommata grandis* | 133 | http://www.nies.go.jp; Ejsmont-Karabin et al. 2004 |
| 110 | *Monommata longiseta* | 96 | http://www.nies.go.jp; Ejsmont-Karabin et al. 2004 |
| 111 | *Monommata phoxa* | 150 | http://www.nies.go.jp; Ejsmont-Karabin et al. 2004 |
| 112 | *Mytilina mucronata* | 191 | http://www.nies.go.jp; Ejsmont-Karabin et al. 2004 |
| 113 | *Notholca acuminata* | 253 | Ejsmont-Karabin et al. 2004 |
| 114 | *Notholca caudata* | 373 | Ejsmont-Karabin et al. 2004 |
| 115 | *Notholca squamula* | 155 | Ejsmont-Karabin et al. 2004 |
| 116 | *Notommata contorta* | 333 | Ejsmont-Karabin et al. 2004 |
| 117 | *Notommata cyrtopus* | 236 | Ejsmont-Karabin et al. 2004 |
| 118 | *Notommata tripus* | 175 | Ejsmont-Karabin et al. 2004 |
| 119 | *Philodina citrina* | 378 | http://www.nies.go.jp |
| 120 | *Philodina megalotrocha* | 203 | http://www.nies.go.jp |
| 121 | *Philodina roseola* | 421 | http://www.nies.go.jp |
| 122 | *Platyias quadricornis* | 260 | Ejsmont-Karabin et al. 2004 |
| 123 | *Pleurotrocha petromyzon* | 270 | http://www.nies.go.jp; Ejsmont-Karabin et al. 2004 |
| 124 | *Ploesoma hudsoni* | 400 | Ejsmont-Karabin et al. 2004 |
| 125 | *Ploesoma triacanthum* | 177 | Ejsmont-Karabin et al. 2004 |
| 126 | *Ploesoma truncatum* | 178 | Ejsmont-Karabin et al. 2004 |
| 127 | *Polyarthra dolichoptera* | 125 | Ejsmont-Karabin et al. 2004 |
| 128 | *Polyarthra euryptera* | 203 | Ejsmont-Karabin et al. 2004 |
| 129 | *Polyarthra longiremis* | 170 | Ejsmont-Karabin et al. 2004 |
| 130 | *Polyarthra major* | 168 | Ejsmont-Karabin et al. 2004 |
| 131 | *Polyarthra minor* | 79 | Ejsmont-Karabin et al. 2004 |
| 132 | *Polyarthra remata* | 96 | Ejsmont-Karabin et al. 2004 |
| 133 | *Polyarthra vulgaris* | 141 | Ejsmont-Karabin et al. 2004 |
| 134 | *Pompholyx sulcata* | 111 | Ejsmont-Karabin et al. 2004 |
| 135 | *Proales decipiens* | 200 | Ejsmont-Karabin et al. 2004 |
| 136 | *Proales fallaciosa* | 260 | Ejsmont-Karabin et al. 2004 |
| 137 | *Proales minima* | 65 | Ejsmont-Karabin et al. 2004 |
| 138 | *Proalinopsis caudatus* | 213 | http://www.nies.go.jp; Ejsmont-Karabin et al. 2004 |
| 139 | *Ptygura crystallina* | 453 | Ejsmont-Karabin et al. 2004 |
| 140 | *Rotaria citrina* | 850 | http://www.nies.go.jp |

| | | | |
|---|---|---|---|
| 141 | *Rotaria elongata* | 1500 | http://www.nies.go.jp |
| 142 | *Rotaria macroceros* | 256 | http://www.nies.go.jp |
| 143 | *Rotaria macrura* | 700 | http://www.nies.go.jp |
| 144 | *Rotaria neptunia* | 573 | http://www.nies.go.jp |
| 145 | *Rotaria rotatoria* | 655 | http://www.nies.go.jp |
| 146 | *Rotaria tardigrada* | 531 | http://www.nies.go.jp |
| 147 | *Rotaria trisecata* | 1150 | http://www.nies.go.jp |
| 148 | *Sinantherina socialis* | 1360 | Ejsmont-Karabin et al. 2004 |
| 149 | *Squatinella rostrum* | 184 | Ejsmont-Karabin et al. 2004 |
| 150 | *Synchaeta lakowitziana* | 285 | http://www.nies.go.jp; Ejsmont-Karabin et al. 2004 |
| 151 | *Synchaeta longipes* | 184 | Ejsmont-Karabin et al. 2004 |
| 152 | *Synchaeta oblonga* | 223 | Ejsmont-Karabin et al. 2004 |
| 153 | *Synchaeta pectinata* | 395 | Ejsmont-Karabin et al. 2004 |
| 154 | *Synchaeta stylata* | 221 | Ejsmont-Karabin et al. 2004 |
| 155 | *Synchaeta tremula* | 226 | Ejsmont-Karabin et al. 2004 |
| 156 | *Synchaeta kitina* | 132 | http://www.nies.go.jp |
| 157 | *Synchaeta grandis* | 500 | Ejsmont-Karabin et al. 2004 |
| 158 | *Taphrocampa annulosa* | 171 | Ejsmont-Karabin et al. 2004 |
| 159 | *Taphrocampa selenura* | 211 | Ejsmont-Karabin et al. 2004 |
| 160 | *Testudinella emarginula* | 114 | http://www.nies.go.jp |
| 161 | *Testudinella parva* | 105 | http://www.nies.go.jp; Ejsmont-Karabin et al. 2004 |
| 162 | *Testudinella patina* | 177 | http://www.nies.go.jp; Ejsmont-Karabin et al. 2004 |
| 163 | *Trichocerca bicristata* | 256 | Ejsmont-Karabin et al. 2004 |
| 164 | *Trichocerca bidens* | 148 | Ejsmont-Karabin et al. 2004 |
| 165 | *Trichocerca birostris* | 173 | http://www.nies.go.jp |
| 166 | *Trichocerca brachyura* | 91 | Ejsmont-Karabin et al. 2004 |
| 167 | *Trichocerca capucina* | 268 | Ejsmont-Karabin et al. 2004 |
| 168 | *Trichocerca cavia* | 113 | Ejsmont-Karabin et al. 2004 |
| 169 | *Trichocerca elongata* | 302 | Ejsmont-Karabin et al. 2004 |
| 170 | *Trichocerca iernis* | 174 | Ejsmont-Karabin et al. 2004 |
| 171 | *Trichocerca intermedia* | 100 | http://www.nies.go.jp |
| 172 | *Trichocerca longiseta* | 261 | Ejsmont-Karabin et al. 2004 |
| 173 | *Trichocerca minuta* | 98 | http://www.nies.go.jp |
| 174 | *Trichocerca porcellus* | 153 | Ejsmont-Karabin et al. 2004 |
| 175 | *Trichocerca pusilla* | 76 | Ejsmont-Karabin et al. 2004 |
| 176 | *Trichocerca rattus* | 172 | Ejsmont-Karabin et al. 2004 |
| 177 | *Trichocerca marina* | 166 | http://www.nies.go.jp |
| 178 | *Trichocerca rotundata* | 112 | http://www.nies.go.jp |
| 179 | *Trichocerca rousseleti* | 88 | Ejsmont-Karabin et al. 2004 |
| 180 | *Trichocerca scipio* | 165 | Ejsmont-Karabin et al. 2004 |
| 181 | *Trichocerca similis* | 155 | Ejsmont-Karabin et al. 2004 |
| 182 | *Trichocerca sulcata* | 110 | Ejsmont-Karabin et al. 2004 |
| 183 | *Trichocerca tenuior* | 153 | Ejsmont-Karabin et al. 2004 |
| 184 | *Trichocerca tigris* | 170 | Ejsmont-Karabin et al. 2004 |
| 185 | *Trichocerca uncinata* | 83 | Ejsmont-Karabin et al. 2004 |
| 186 | *Trichocerca weberi* | 112 | Ejsmont-Karabin et al. 2004 |
| 187 | *Trichotria pocillum* | 127 | http://www.nies.go.jp |
| 188 | *Trichotria tetractis* | 131 | http://www.nies.go.jp |
